# Nascent transcription profiling identifies H3K36 methylation as a custodian of the somatic transcriptome

**DOI:** 10.64898/2026.10.25.761580

**Authors:** Hina Sultana, Vasudha Vandadi, Harmony R. Salzler, Apoorva Baluapuri, Darshan Ponkiya, Alexis K. Hirth, Grace C. Vernesoni, Seth R. Goldman, Karen Adelman, A. Gregory Matera

**Affiliations:** Integrative Program for Biological and Genome Sciences, University of North Carolina, Chapel Hill, NC, USA; Department of Biological Chemistry and Molecular Pharmacology, Blavatnik Institute, Harvard Medical School, Boston, MA, USA; Department of Biology, University of North Carolina, Chapel Hill, NC, USA; Department of Genetics and Lineberger Comprehensive Cancer Center, University of North Carolina, Chapel Hill, NC, USA

## Abstract

Faithful gene expression not only requires activation of productive transcription units, but also suppression of aberrant initiation within or between gene bodies. In *Saccharomyces*, Set2-mediated H3 lysine-36 methylation (H3K36me) recruits a deacetylase complex that removes acetylation deposited behind elongating RNA polymerase II, preventing cryptic initiation. Whether metazoans require Set2 or the K36 residue itself for this function has been difficult to test. Here, we combined PRO-seq, RNA-seq, and chromatin profiling in *Drosophila* wing imaginal discs to define individual contributions of Set2, H3.2K36 and H3.3K36 to transcriptome fidelity. Intragenic initiation events are abundant but largely unresponsive to loss of K36 methylation. Aberrant initiation instead concentrates in repetitive, extragenic chromatin, co-marked by H3K9me2/3 and H3K36me2 but depleted of H3K36me3, a dual heterochromatin signature that is also conserved in mammals. Within this compartment, piRNA clusters, germline-restricted genes, and LTR and LINE retrotransposons are aberrantly activated in somatic tissue. A K36R substitution in *H3.2*, but not in *H3.3*, derepresses these loci to a far greater extent than deleting *Set2*, which retains K36me2 but cannot deposit K36me3. Thus, metazoan H3K36 safeguards the somatic transcriptome principally within heterochromatin, rather than within gene bodies.

## Introduction

Accurate transcription initiation is fundamental to cellular identity and genome integrity. For the purposes of this study, we define transcriptome fidelity as the degree to which the proper complement of RNA species is produced from a given genome. Eukaryotic chromatin must permit productive elongation while suppressing initiation at sites that are not *bona fide* promoters. Set2 associates with the phosphorylated C-terminal domain of elongating RNA polymerase II (Pol II CTD) and deposits methylation on histone H3 lysine-36 (H3K36me) across transcribed regions, a coupling that is conserved from yeast to humans (Krogan et al., 2003; Xiao et al., 2003; Yoh et al., 2008). The function of this residue was first characterized in budding yeast, wherein H3K36me3 was shown to recruit a deacetylase complex (Rpd3S) that erases promoter-like acetylation and prevents cryptic transcription within gene bodies (Carrozza et al., 2005; Joshi and Struhl, 2005; Keogh et al., 2005). Whether this anti-cryptic function is a general eukaryotic solution, or a specialization suited to the yeast genome, remains an open question.

These early studies established that loss of Set2, or mutation of H3K36, triggers pervasive cryptic transcription within coding regions, firmly linking H3K36me3 to suppression of intragenic initiation. Three features of this model are specific to *Saccharomyces.* Animal genomes contain two types of H3 histones, a replication-independent H3.3 and replication-dependent H3.2/H3.1, whereas budding yeast has but a single H3.3 ortholog (Ahmad and Henikoff, 2002). Further, Set2 is the only budding yeast H3K36 methyltransferase, whereas metazoans encode several, including the NSD and Ash1 enzymes that deposit H3K36me1/2 independently of elongating Pol II (Bell et al., 2007). As such, H3K36me3 is deposited and read within transcription units, which occupy most of the yeast genome but a minority of an animal one. In contrast, the downstream machinery is conserved: *Drosophila* MRG15, for example, is the Eaf3 orthologue and functions within an analogous metazoan complex (Zhang et al., 2006).

Analyses in animal cells have yielded mixed results. In *Drosophila*, replacement of *H3.2* genes with *H3.2^K36R^* missense alleles was reported to cause global hyperacetylation of histone H4, yet intragenic cryptic initiation events were comparatively rare and thus may be cell-type or context dependent (Meers et al., 2017). RNAi-mediated depletion of human SETD2 was reported to alter intragenic initiation in only ∼11% of active genes, but the extent and functional significance of these events remain controversial (Carvalho et al., 2013). This apparent disconnect between hyperacetylation and cryptic initiation in metazoans could indicate that surveillance and removal of cryptic transcripts is more efficient, or that compensatory mechanisms exist. A third possibility is that the principal requirement for H3K36 lies in genomic compartments that budding yeast does not possess.

Animal genomes carry large intergenic (extragenic) regions, occupied in substantial part by transposons and other repetitive sequences, and organized into extensive domains of constitutive heterochromatin. The budding yeast genome has few of these features, and lacks H3K9 methylation, HP1, and RNAi, systems that *S. pombe* retains (Grewal and Jia, 2007). These are the compartments in which a metazoan genome must suppress inappropriate transcription (Brennecke et al., 2007; Grewal and Elgin, 2007; Slotkin and Martienssen, 2007; Sienski et al., 2012), and the canonical anti-cryptic initiation model comes from an organism that lacks them.

Whether H3K36 methylation is required in those compartments has been difficult to test, for three reasons. First, metazoans possess multiple histone H3 variants that are differentially distributed across the genome and may respond differently to methylation. Second, deleting or depleting Set2/SETD2 removes not only the catalytic activity, but also potential structural roles these enzymes may play in transcription complexes (Ambrosi et al., 2026). More importantly, it does not remove H3K36 methylation. It removes a single writer, and leaves H3K36me marks deposited by NSD and Ash1 intact. Clearly, enzyme deletion and residue substitution are not equivalent perturbations. Removing Set2 tests the requirement for its trimethyl product and/or roles in other processes, whereas removing the residue tests the requirement for K36 in any methyl state. Third, most studies have relied on whole-organism or heterogeneous cell population analyses, which may obscure tissue-specific cryptic transcription events.

Our own previous work illustrates the problem. Start-seq analysis of animals carrying an *H3.2^K36R^* substitution identified over 17,000 novel unannotated transcription start sites (nuTSSs). The differentially expressed ones were predominantly extragenic, and those mapping within genes showed no enrichment in mutants relative to controls (Meers et al., 2017). That study, however, was subject to all three limitations: it was restricted to replication-dependent H3, included no *Set2* allele for comparison, and was performed on mixed tissues (whole larvae).

Here, we address these limitations by integrating high-resolution nascent-transcription profiling (PRO-seq), steady-state RNA expression analysis (polyA-selected RNA-seq), and chromatin profiling (CUT&RUN) to define how Set2 and H3K36me maintain transcriptome fidelity. The experiments were carried out in wandering third-instar wing imaginal disc, a tissue composed of relatively homogeneous populations of diploid epithelial cells that give rise to the adult wing, and free from the confounding heterogeneity of other, largely polyploid, larval tissues. Furthermore, because all replication-dependent histone genes in *Drosophila* reside at a single locus (*HisC*), we employed histone gene replacement strategies (McKay et al., 2015; Meers et al., 2018a) to remove the methyl-accepting residue itself. We examined three matched mutant and control pairs, each in biological triplicate: *Set2*Δ against *yw*, replication-dependent H3.2^K36R^ against its wild-type transgenic control, and replication-independent *H3.3^K36R^* against its control.

We find that initiation events within gene bodies are abundant in all genotypes, but those that respond to loss of H3K36me are not concentrated there, and the sites that do respond fail to correlate with hyperacetylation of H4. Aberrant initiation is instead more pronounced in repetitive, extragenic regions of the genome. These derepressed loci largely reside in regions that are co-marked by H3K9me2/3 and H3K36me2. Steady-state transcriptome profiling (RNA-seq) illustrates the consequence: piRNA clusters and germline-restricted genes, including piRNP biogenesis factors, are aberrantly activated in the soma, alongside LTR and LINE retrotransposons. Within this dual heterochromatin (dual HC) compartment, *H3.2^K36R^* mutants activate cryptic loci far more frequently than *Set2*Δ mutants, which retain K36me2 but cannot deposit K36me3, whereas *H3.3^K36R^* mutants show no such activation. H3K36me2 is therefore required for silencing in this compartment. Together, these results identify H3K36 methylation as a custodian of the somatic transcriptome, and place its custodial function within heterochromatin rather than within gene bodies.

## Results

### PRO-seq defines the transcription start site landscape of the wing imaginal disc

To determine how Set2 and H3K36 methylation maintain transcriptome fidelity in a single tissue, we assembled six genotypes as three matched mutant and control pairs and processed them in parallel (Fig. 1A; Fig. S1). From wing imaginal discs of wandering third-instar larvae, we prepared nuclei for PRO-seq, RNA for polyA-selected RNA-seq, and chromatin for CUT&RUN targeting acetylated histone H4 (Fig. S2A-F).

**Figure 1.**
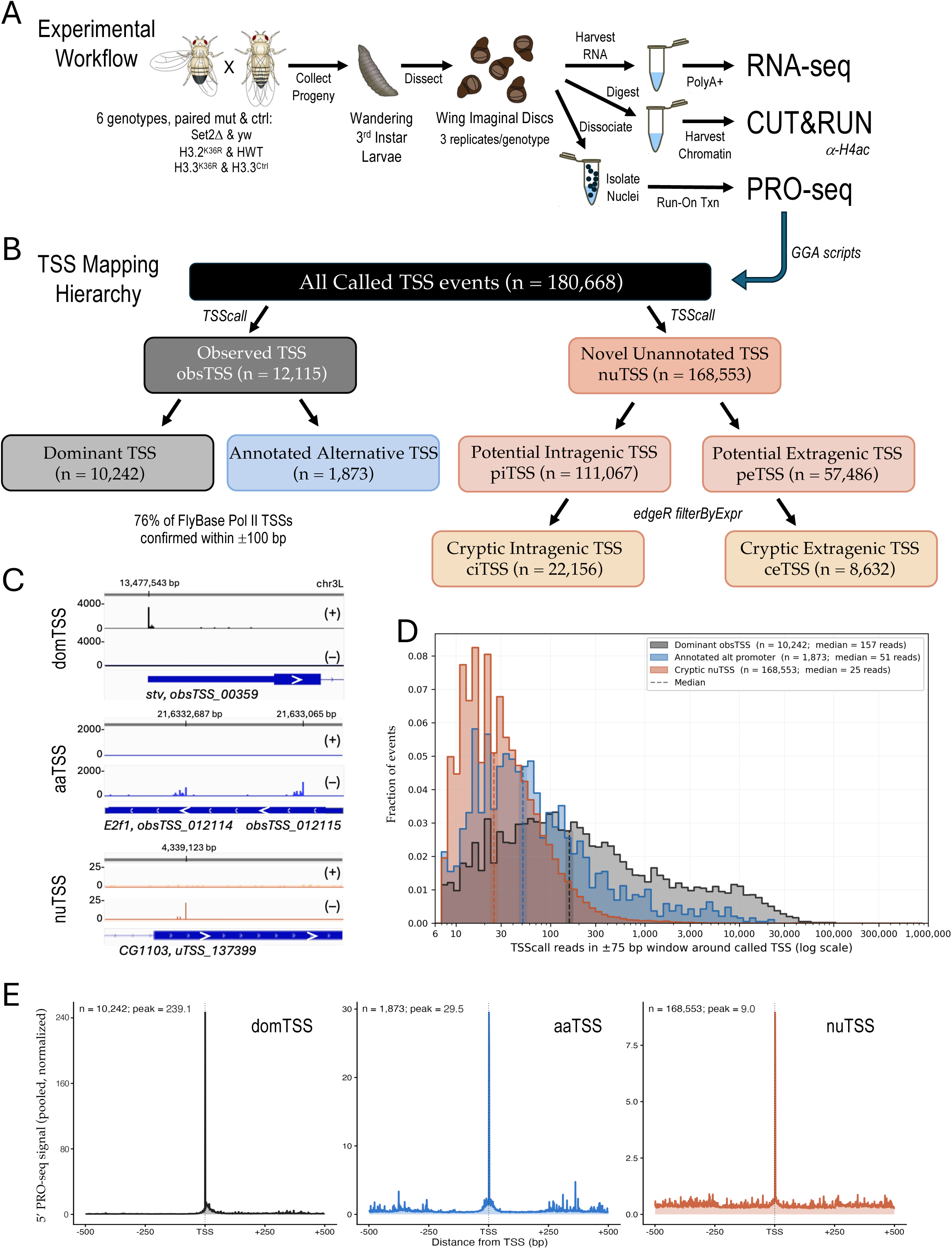
PRO-seq defines the transcription start site landscape of the wing imaginal disc. **(A)** Experimental design. Wing imaginal discs from wandering third-instar larvae of three matched mutant and control pairs (*Set2*Δ and *yw*; *H3.2^K36R^* and *H3.2^HWT^*; *H3.3^K36R^* and *H3.3^Ctrl^*) provided nuclei for PRO-seq, RNA for polyA-selected RNA-seq, and chromatin for CUT&RUN against pan-acetylated histone H4. **(B)** Hierarchy of transcription start site classification, from calls made on pooled data from all six genotypes. obsTSS, observed TSS; domTSS, dominant TSS; aaTSS, annotated alternative TSS; nuTSS, novel unannotated TSS; piTSS and peTSS, potential intragenic and extragenic TSSs; ciTSS and ceTSS, the subsets retained for differential analysis. **(C)** Pooled 5′ PRO-seq signal at *stv* (a domTSS), *E2f1* (two aaTSSs) and *u_TSS137399* (a nuTSS initiating antisense to its host gene, *CG1103*). **(D)** Distribution of PRO-seq signal by class, normalized to sum to 1 within each class. Dashed lines indicate median read depth. **(E)** Metaplots of aggregate 5′ PRO-seq profiles. Note that the three panels use different y-axis scales.

Nascent transcription was profiled by PRO-seq, wherein transcriptionally engaged polymerases incorporate a biotinylated nucleotide at the 3′ end of nascent RNAs (Kwak et al., 2013; Mahat et al., 2016). Sequencing the paired ends within these libraries recovers the 5′ end of each read. Because fragmentation and degradation also generate 5′ ends, a single read is not evidence of initiation. We therefore called transcription start sites (TSSs) as positions upon which the 5′ ends of multiple independent reads coincide, using TSScall (Scruggs et al., 2015). Calls were made once on pooled data from all six genotypes, so that the same set of sites is tested in every comparison.

TSSs were first classified into two groups: those that had been previously observed and those that had not (Fig. 1B). Of 180,668 called events, 12,115 fell within ±100 bp of an annotated start in FlyBase and were designated observed TSSs (obsTSS). These sites belong to 10,242 genes active in the wing disc, including snRNA and snoRNA as well as mRNA genes. For each active gene, we designated the most heavily used site as its dominant TSS (domTSS). Roughly 18% of active genes (1,873/10,242) initiated from more than one site. These were termed annotated alternative starts (aaTSSs). The remaining 168,553 calls were designated novel unannotated sites (nuTSSs) and partitioned into 111,067 potential intragenic (piTSS) and 57,486 potential extragenic (peTSS) events. Overall, this analysis recovered 76% of annotated Pol II TSSs to within ±100 bp. Note, this is a lower bound because the FlyBase annotation aggregates start sites across all tissues and developmental stages.

Sites in both classes satisfy the read convergence criterion and represent genuine initiation events (Nechaev et al., 2010; Meers et al., 2018b). Many nuTSSs, however, were detected in a single replicate. Filtering for expression (edgeR filterByExpr; Methods) retained 22,156 cryptic intragenic TSSs (ciTSS) and 8,632 cryptic extragenic TSSs (ceTSS) for differential analysis, quantified in a 21 bp window, as 94% of *Drosophila* TSSs are narrower than 20 bp (Meers et al., 2018b). Individual loci illustrate the sharpness of fly promoters (Fig. 1C). At *stv*, the domTSS accounts for essentially all initiation across the gene; at *E2f1*, two annotated alternative starts are used simultaneously. Within *CG1103*, an unannotated antisense site is two orders of magnitude weaker than the domTSS at *stv*, but its peak is equally well defined.

Signal distributions differ by class but overlap substantially (Fig. 1D), and the aggregate profiles show the same ordering (Fig. 1E). Most of the TSSs in the wing dataset lie outside the current annotation, and two-thirds of those fall within gene bodies, the compartment in which the yeast model predicts that H3K36 methylation is required.

### Cryptic intragenic initiation is uncoupled from host gene activity and H4 acetylation

To determine whether loss of H3K36 methylation activates cryptic start sites within gene bodies, we compared 5′ PRO-seq signal between each mutant and its matched control at all 22,156 ciTSSs (Fig. 2A). Each site was treated as an independent count feature, so the analysis reports whether initiation at that position changes, rather than whether output from the surrounding gene changes. Activation was detectable but rare: 291 sites were significantly upregulated in *Set2*Δ, 222 in *H3.2^K36R^*, and 47 in *H3.3^K36R^*, between 0.2% and 1.3% of those tested.

**Figure 2.**
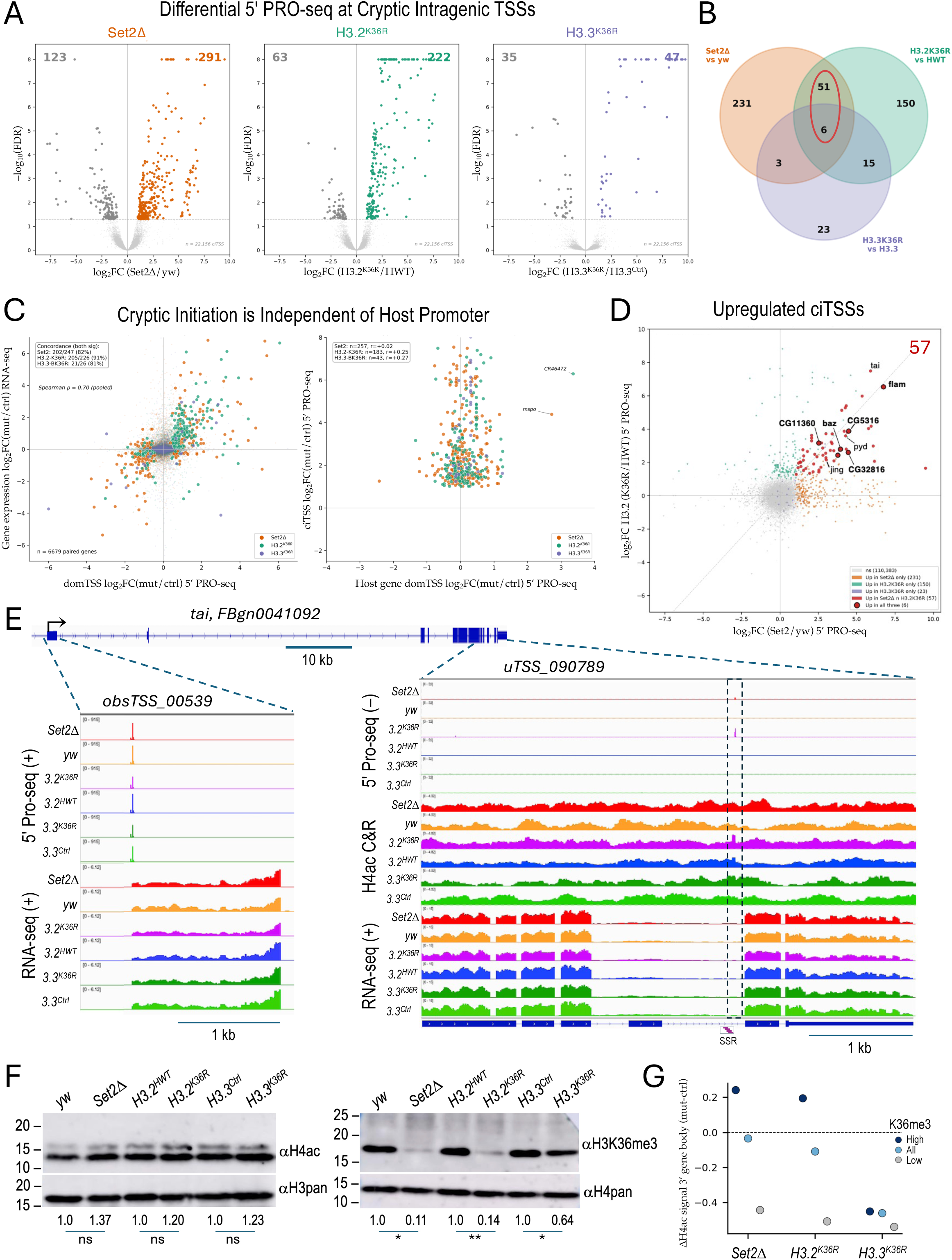
Cryptic intragenic initiation is uncoupled from host gene activity and histone H4 acetylation. **(A)** Differential 5′ PRO-seq signal at the 22,156 ciTSSs, shown as volcano plots for each mutant against its matched control (edgeR; FDR < 0.05 and |log2FC| > 1). **(B)** Overlap of upregulated ciTSSs between genotypes. Circled region indicates the 57 sites shared between *Set2*Δ and *H3.2^K36R^*. **(C)** Left: log₂ fold change in domTSS 5′ PRO-seq signal against log₂ fold change in polyA RNA-seq, for 6,679 genes passing threshold in both assays. Inset gives directional concordance among genes significant in both assays. Right: log₂ fold change at upregulated ciTSSs against log₂ fold change at the domTSS of the same host gene. **(D)** Scatter plot of log2FC 5’ PRO-seq values for *Set2*Δ and *H3.2^K36R^* at all piTSSs (gray). The 57 shared sites identified in panel B are shaded in red; other colored points are as noted in the key. **(E)** H4ac CUT&RUN, 5′ PRO-seq and RNA-seq at the *taiman (tai)* locus: promoter region (left) and the cryptic intragenic site within the dashed box (right). SSR, simple sequence repeat. **(F)** Western blots of whole-larval lysates from all six genotypes: anti-acetylated H4 with anti-panH3 loading control (left), and anti-H3K36me3 with anti-panH4 loading control (right). Values below each lane give signal relative to the matched control; quantification is shown in Fig. S2G. (G) Mean difference (mutant − control) in H4ac CUT&RUN signal over the 3′ half of the gene body, for genes with high K36me3 (dark blue), all expressed genes (light blue) and genes with low K36me3 (gray). Full profiles are shown in Fig. S2H,I.

The two genotypes that lose K36 methylation from replication-dependent histones converge on a small common set: 57 upregulated ciTSSs are shared between *Set2*Δ and *H3.2^K36R^* (Fig. 2B). *H3.3^K36R^* contributes only 23 sites of its own, and this weak response of the replication-independent variant recurs throughout the analyses below.

If cryptic initiation within a gene body were licensed by loss of the mark deposited during that gene’s own transcription, activation should mirror the activity of the host promoter. It does not (Fig. 2C). Across 6,679 genes, domTSS output correlated strongly with mRNA level (pooled Spearman ρ = 0.70), and genes significant in both assays agreed in direction in 81– 91% of cases. The assay therefore detects coupling where coupling exists.

Cryptic intragenic sites behave differently. They rose as much as 90-fold, whereas the domTSS of the same host gene changed little, and the two were only weakly correlated (r = 0.02 to 0.27). Comparing effect sizes between genotypes identifies the loci that respond most strongly (Fig. 2D), including cryptic sites within *taiman* (*tai*) and *CG5316*. Of the eight highlighted loci, only the *CG5316* site lies in uniquely mappable sequence; the *tai* site directly abuts a simple sequence repeat (Fig. 2E), and the remainder fall directly within transposons.

The yeast model further predicts that cryptic sites should be locally hyperacetylated. In bulk chromatin, H4ac was not significantly changed in any mutant relative to its matched control, whereas H3K36me3 on the same extracts fell to 0.11, 0.14 and 0.64 of control (Figs. 2F; S2G). PRO-seq spike-in recovery was likewise consistent across genotypes, and CUT&RUN normalization factors (TMM) were approximately 1.0 in every comparison (Methods). Bulk acetylation is thus unchanged, but its distribution along genes is not. In *Set2Δ* and *H3.2^K36R^*, H4ac rises over the 3′ halves of genes that carry high K36me3 and falls over genes with little K36me3, whereas *H3.3^K36R^* shows no such dependence (Fig. 2G; Fig. S2H,I).

Local quantification is constrained by sequence. Although only 6.8% of the 22,156 ciTSSs overlap an annotated repeat, 36.8%, 63.1% and 87.2% of the upregulated sites in *Set2*Δ, *H3.2^K36R^* and *H3.3^K36R^* do, and the ciTSS clusters at *pyd* and *jing* lie within transposons. Short-read CUT&RUN cannot be reliably quantified in such regions, so we restrict quantitative statements to repeat classes rather than to specific locations.

Three features of these data are difficult to reconcile with a Set2•Rpd3S suppression model operating in this tissue. Cryptic initiation within gene bodies is rare, it is uncoupled from the transcription of the host gene, and it occurs without detectable changes in H4 acetylation. Hence, whatever H3K36 methylation is doing to maintain metazoan transcriptome fidelity, it is not principally functioning inside gene boundaries.

### Cryptic extragenic initiation is widespread and concentrated in H3K9-methylated chromatin

If the requirement for H3K36 methylation lies outside gene bodies, cryptic extragenic sites should respond where the intragenic ones did not. We therefore applied the same analysis to the 8,632 ceTSSs (Fig. 3A). The response was substantial: 1,579 sites were significantly upregulated in *Set2*Δ, 1,887 in *H3.2^K36R^* and 638 in *H3.3^K36R^*. Downregulated sites occurred at a constant fraction (∼40%) of the upregulated frequency in every genotype, defining a symmetric baseline of stochastic change against which the upregulated signal stands out. Upregulated sites account for 18.3%, 21.9% and 7.4% of tested extragenic sites, against 1.31%, 1.00% and 0.21% of intragenic ones; cryptic activation is thus 14- to 35-fold greater outside genes than within them.

**Figure 3.**
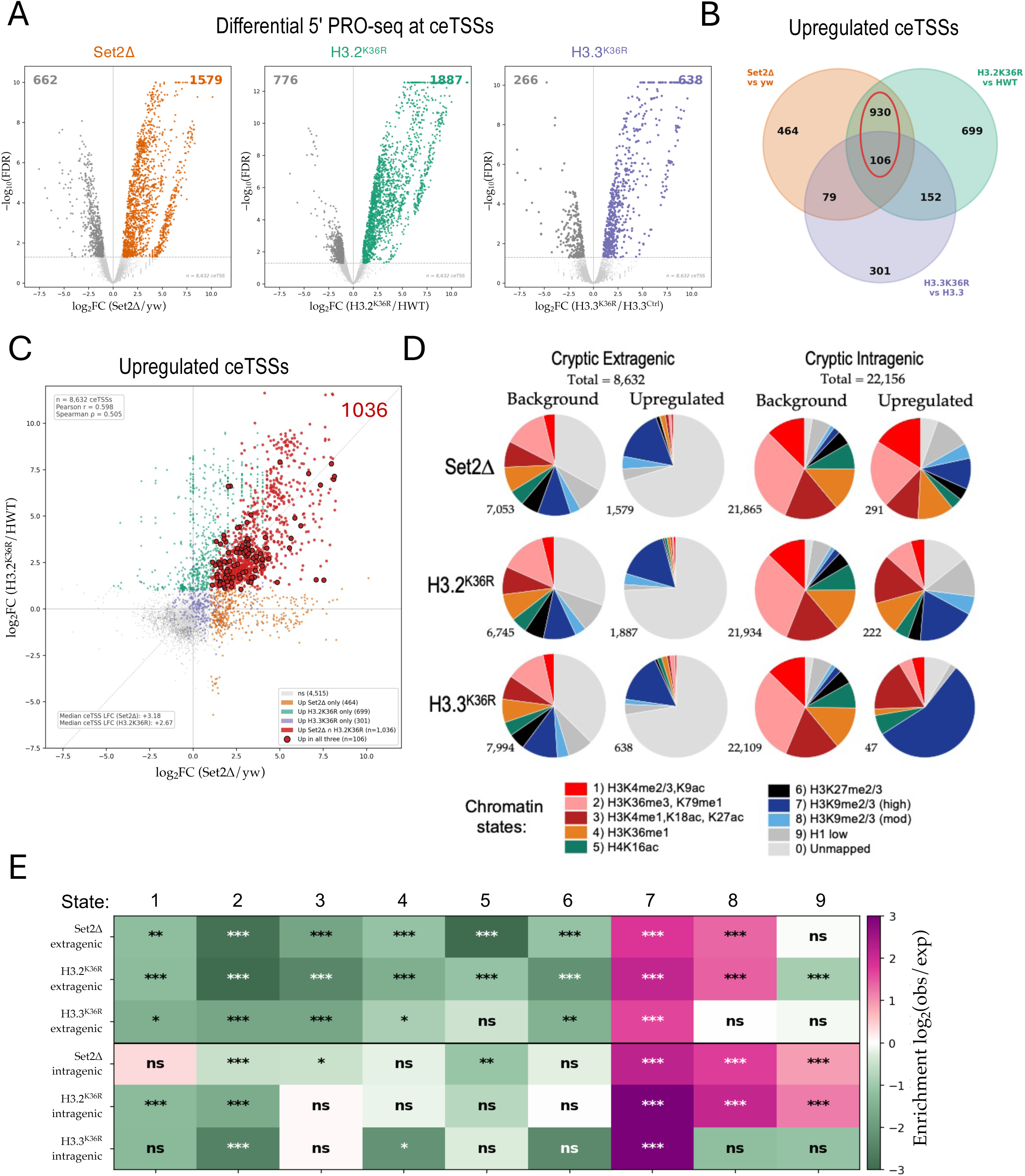
Cryptic extragenic initiation is widespread and concentrated in H3K9-methylated chromatin. **(A)** Differential 5′ PRO-seq signal at the 8,632 ceTSSs, analyzed as in Figure 2A. **(B)** Venn diagram of upregulated ceTSSs between genotypes. **(C)** Pairwise comparison of ceTSS effect sizes between *Set2*Δ and *H3.2K36R*, across all tested sites. Point categories shaded as keyed. **(D)** Chromatin state composition of upregulated ceTSSs and of the tested background, by genotype, using the nine-state BG3 model (Kharchenko et al., 2011). Unmapped sites (state 0) are included. **(E)** Enrichment of upregulated sites over the tested background of the same class, expressed as log₂(observed/expected), for extragenic (upper three rows) and intragenic (lower three rows) cryptic TSSs, among state-assigned sites only. Two-sided Fisher’s exact test, uncorrected: * p < 0.05, ** p < 0.01, *** p < 0.001; ns, not significant.

Unlike the intragenic ones, the extragenic responses converged: 1,036 upregulated ceTSSs are shared between *Set2*Δ and *H3.2^K36R^*, and 106 are shared across all three genotypes (Fig. 3B). Effect sizes in the two genotypes also correlate across the full tested set (Pearson r = 0.60; Fig. 3C), whereas the same comparison at intragenic sites showed almost no relationship (Fig. 2D). Thus, enzyme loss and residue loss are distinguishable within gene bodies but not outside of them.

We next classified the responsive sites using the nine-state chromatin model derived from BG3 cells (Kharchenko et al., 2011). The state composition of upregulated ceTSSs differs from that of the tested background (Fig. 3D), and enrichment over expectation identifies the states responsible (Fig. 3E). Upregulated ceTSSs are enriched in state 7 (high H3K9me2/3) in all three genotypes and in state 8 (moderate H3K9me2/3) in *Set2*Δ and *H3.2^K36R^*, and are depleted from the active states (states 1–5). Cryptic extragenic initiation is thus concentrated within H3K9me chromatin.

The *H3.3^K36R^* response is narrower, concentrated in state 7 and in the unmapped repeat fraction, consistent with the known enrichment of H3.3 in pericentromeric and repeat-associated heterochromatin. Remarkably, the same signature appears within gene bodies: upregulated ciTSSs are also enriched in state 7 in all three genotypes, and in state 8 in *Set2*Δ and *H3.2^K36R^* (Fig. 3E, lower rows). The intragenic sites that do respond are therefore the heterochromatic minority, and they share a chromatin environment with the extragenic sites rather than with the gene bodies that surround them. A site’s position relative to an annotated gene predicts the response only weakly, whereas H3K9-methylated chromatin predicts it in both compartments.

### Cryptic initiation concentrates at transposons and piRNA clusters, including germline-restricted loci

To locate the responsive sites, we plotted all 30,788 cryptic intragenic and extragenic TSSs by genomic coordinate against their pooled mean log2 fold change (Fig. 4A). Along each chromosome arm, sites change modestly in both directions, reproducing at genome scale the symmetric baseline noted above. Against that background, the large fold-changes are almost exclusively increases. They fall in the pericentromeric region of every major arm, across chromosomes 4 and Y, and in a discrete block on the X chromosome, and the sites shared between genotypes coincide with these same positions.

**Figure 4.**
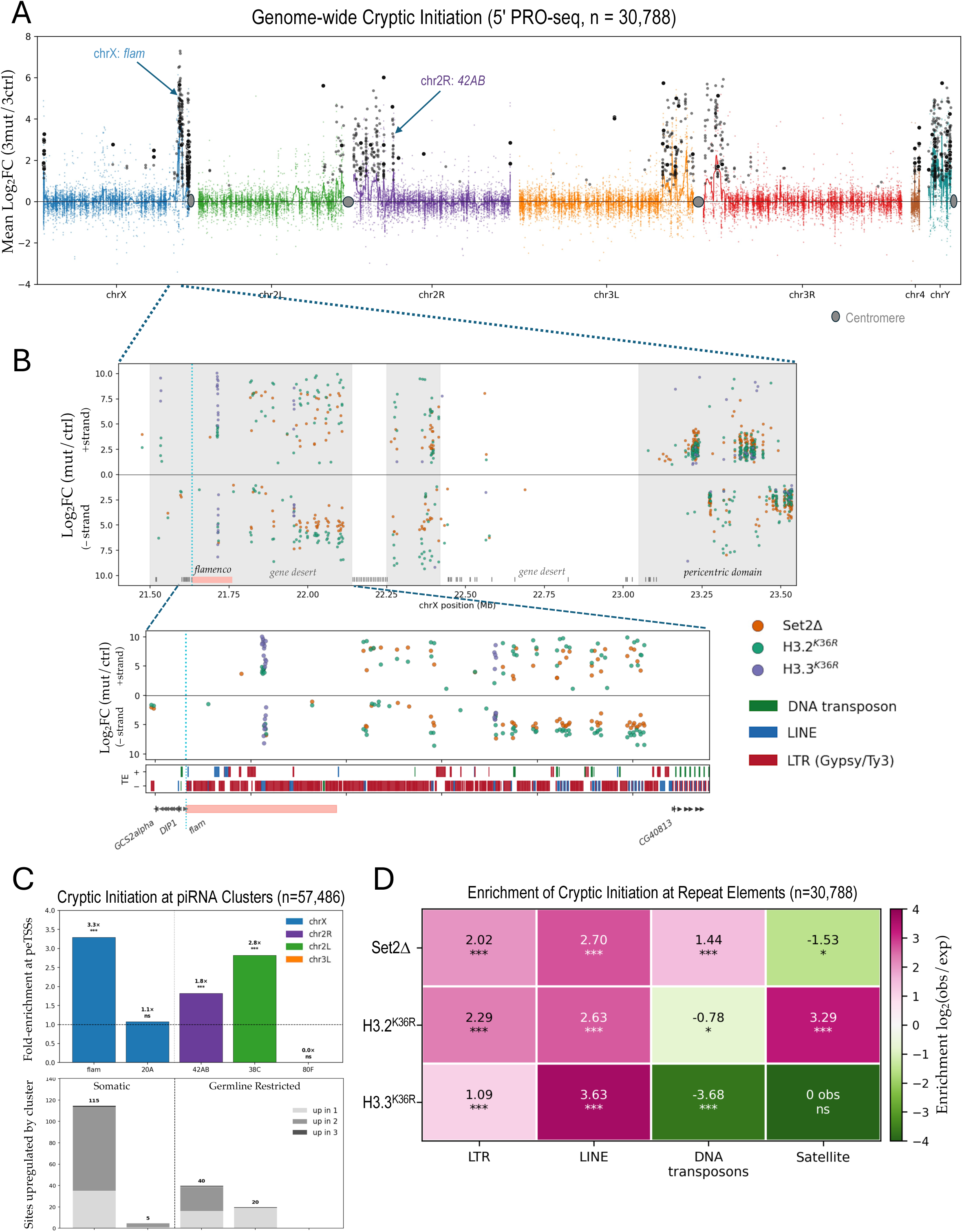
Cryptic initiation concentrates at transposons and piRNA clusters, including germline-restricted loci. (**A**) Manhattan plot of all 30,788 cryptic intragenic and extragenic TSSs, plotted by chromosomal coordinate against the pooled mean log₂ fold change across the three mutant/control comparisons, and colored by chromosome arm. Sites upregulated in both *Set2*Δ and *H3.2^K36R^* (Fig. 3B) are enlarged and shaded in dark gray, or in black if up in all three genotypes. (**B**) The centromere-proximal region of the X chromosome containing the *flamenco (flam)* locus (chrX:21.4–23.6 Mb). Top, log2 fold change of significantly upregulated sites for each mutant, with plus-strand (+) sites above and minus-strand (–) sites below. Bottom, the gene desert proximal to *flam*; the *flam* domTSS is unchanged in all three genotypes. Tracks show gene models and transposons by class and by orientation relative to *flam*. (**C**) Enrichment of upregulated extragenic TSSs at five piRNA clusters, against the full set of 57,486 peTSSs (Fisher’s exact test; *** p < 0.001; ns, not significant). Lower, pooled counts of upregulated sites per cluster, colored by reproducibility across genotypes. (**D**) Enrichment of upregulated cryptic TSSs (intragenic and extragenic pooled) at repeat classes, as log₂(observed/expected). Two-sided Fisher’s exact test with Benjamini-Hochberg correction: *** q < 0.001, ** q < 0.01, * q < 0.05; ns, not significant.

The block on the X chromosome contains *flamenco (flam)*, the major somatic piRNA cluster (Fig. 4B). Across chrX 21.4 to 23.6 Mb, 945 sites are significantly upregulated in at least one genotype, and they resolve into discrete heterochromatic blocks separated by intervals that contain none.

The intervals are as informative as the blocks. Between the two transposon-dense deserts flanking the annotated cluster (∼22.15 to 22.25 Mb) lies an island of 19 genes beginning at *CG40813*, and distal to it lie two other gene-rich intervals; none contains a single upregulated cryptic site in any genotype. These intervals are free of annotated repeats, so the absence of signal is a direct observation rather than a detection failure. Cryptic initiation therefore has sharp boundaries that coincide with the edges of repeat-dense chromatin rather than with gene boundaries or transcriptional status.

The *flamenco* locus is a uni-strand piRNA cluster, transcribed from a single promoter in one orientation, with the transposon copies it targets inserted predominantly in the opposite orientation (Brennecke et al., 2007). Maintaining that arrangement requires initiation to stay confined to one strand. In the derepressed desert proximal to *flamenco* (Fig. 4B, lower), upregulated sites appear on both strands, downstream of the *flamenco* promoter, whose TSS is unchanged in all three genotypes.

Mapping initiation at other known piRNA clusters shows that the response is not restricted to the somatic ones (Fig. 4C). Against all 57,486 detected extragenic sites, *flamenco* is enriched 3.3-fold, and the germline-restricted dual-strand clusters 42AB and 38C 1.8-fold and 2.8-fold, respectively. The somatic locus at 20A is not enriched, and 80F carries no upregulated sites. Both major germline dual-strand clusters are therefore perturbed in a somatic tissue. Because these regions are repeat-dense, we treat these values as a ranking of relative magnitude across clusters rather than as precise fold-enrichments.

Sites in all genotypes fall preferentially on transposons, predominantly LTR and LINE elements (Fig. 4D). Beyond that shared preference, the genotypes differ. Satellite sequence is enriched among *H3.2^K36R^* sites but depleted in *Set2*Δ and absent from the *H3.3^K36R^* response, LINE enrichment is greatest in *H3.3^K36R^*, and *Set2*Δ is the only genotype enriched at DNA transposons.

These assignments were made at the level of repeat class, because multi-mapping reads are assigned at random among equally scoring positions, so copies of a given element return near-identical sets of called sites. Detection also rises with element length, from under 1% of elements below 200 bp to 16% of those above 3 kb, so enrichment for long families such as LINEs partly reflects where a site can be resolved at all. Only 21%, 13% and 5% of upregulated sites in *Set2*Δ, *H3.2^K36R^* and *H3.3^K36R^* lie outside any annotated repeat, so the *H3.3^K36R^* response largely rests on sequences that cannot be individually distinguished. The *flamenco* coordinates are also provisional: the locus varies between fly strains, and long-read assemblies place it well beyond its annotated extent in the dm6 build (Handler and Brennecke, 2026; Moritoh et al., 2026).

### Loss of H3K36me activates a germline transcriptional program in somatic tissue

We reasoned that aberrant initiation in heterochromatin should have consequences for the steady-state transcriptome, so we carried out polyA-selected RNA-seq on the same tissue. The response was biased toward activation in all three comparisons (Fig. 5A; see also Fig. S3). In *Set2*Δ, 482 genes were significantly upregulated and 169 downregulated; in *H3.2^K36R^*, 391 and 83; in *H3.3^K36R^*, 143 and 73. The replication-independent variant again showed the weakest response.

**Figure 5.**
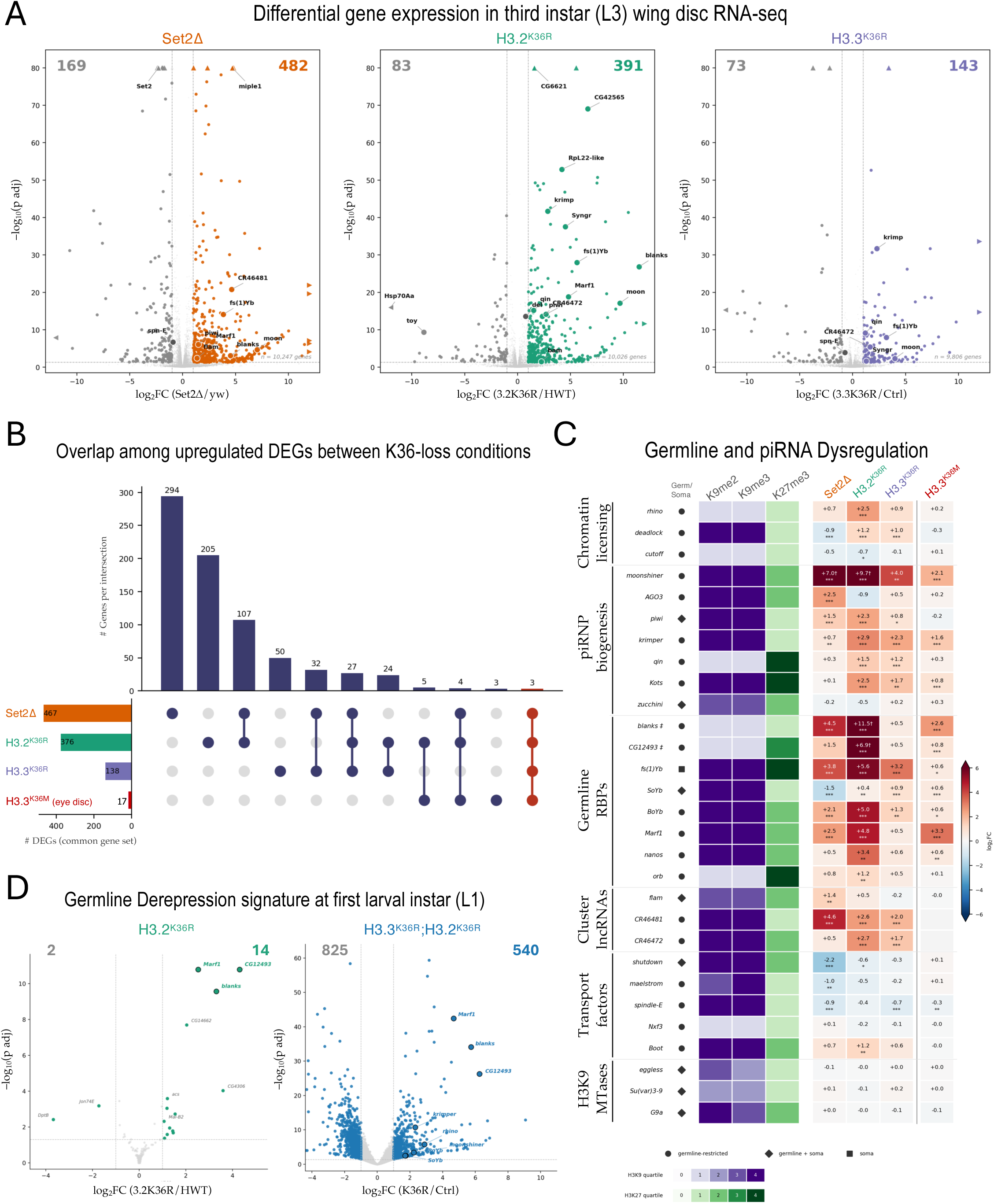
Loss of H3K36me activates a germline transcriptional program in somatic tissue. **(A)** Volcano plots showing differential expression (polyA RNA-seq) for each mutant against its matched control. Labeled points include the germline and piRNA-pathway genes shown in panel C. **(B)** Intersection of upregulated genes across the three wing genotypes and an *H3.3^K36M^* eye-antennal disc dataset (Chaouch et al., 2021), among the 11,080 genes tested in every comparison; red, the three genes shared by all four conditions. **(C)** Log₂ fold change for 29 germline and piRNA-pathway genes, grouped by pathway, with H3K9 methyltransferases as a negative control; H3K9me2/3 and H3K27me3 occupancy quartiles and germline/soma restriction are annotated at left. DESeq2 Wald test: * p < 0.05, ** p < 0.01, *** p < 0.001. **(D)** Differential expression at the first larval instar (L1) in *H3.2^K36R^* (left) and in an *H3.3^K36R^;H3.2^K36R^* combined mutant (right).

Most upregulated genes were restricted to a single genotype, but the three wing genotypes converged on a shared core (Fig. 5B). Among the 11,080 genes tested in every dataset, 107 were upregulated in both *Set2*Δ and *H3.2^K36R^* and 27 in all three wing genotypes. Three genes, *moonshiner*, *CR43961* and *Ir40a*, were also upregulated in eye-antennal discs expressing *H3.3^K36M^* (Chaouch et al., 2021). That set is small (17 genes) because the dominant-negative oncohistone is expressed in only a subset of cells on a wild-type background.

The genes activated in common are recognizably germline ones (Fig. 5C; see also Lindehell et al., 2023). Among 29 germline and piRNA-pathway genes, assigned to tissue-restriction categories independently of the present data, derepression spans functionally distinct arms of the pathway. Among piRNP assembly and biogenesis factors, *moonshiner* is activated in all four conditions, with *krimper*, *AGO3* and *piwi* also elevated. Germline RNA-binding proteins show the largest individual changes: *blanks* rises 11.5 log2 units in *H3.2^K36R^*, and *Marf1*, *fs(1)Yb* and *CG12493* are upregulated across genotypes. The cluster-derived lncRNAs *CR46481* and *CR46472* are activated in all three wing genotypes, whereas the chromatin licensing factors respond least consistently.

This derepression is unlikely to be a secondary consequence of losing the silencing enzymes: the H3K9 methyltransferases *eggless*, *Su(var)3-9* and *G9a* are unchanged in every genotype. Nor does the pathway respond as a coordinated unit, as *shutdown* and *maelstrom* decrease in *Set2*Δ, and *spindle-E* in *Set2*Δ and H3.3^K36R^. The affected genes appear to be escaping repression individually. 5′ PRO-seq confirms this reflects new initiation rather than post-transcriptional accumulation: nascent signal at the domTSS rises specifically in *H3.2^K36R^*, and only among genes silent in the matched control.

Germline genes are among the first loci to respond (Fig. 5D). At the first larval instar, the *H3.2^K36R^* transcriptome is almost unchanged (14 genes up and 2 down of 11,665 tested), yet three of the 14, *CG12493*, *blanks* and *Marf1*, are germline pathway genes, a hundredfold enrichment over expectation (p = 3.1×10⁻⁶). Animals carrying K36R substitutions in both replication-dependent and replication-independent histones show a far broader response (540 genes up, 825 down), within which the same three genes are joined by *rhino*, *krimper*, *moonshiner*, *BoYb* and *SoYb* (7-fold enrichment, p = 1.1×10⁻⁵). Germline derepression thus begins early in larval development and is sharpest where the overall transcriptional disturbance is least.

### Loss of NSD-dependent H3K36me2 triggers cryptic initiation in dual K9me•K36me2 heterochromatin

H3K9 methylation is the central mark of heterochromatic silencing (Schotta et al., 2002), and it also confines lineage-restricted genes, including germline genes, to their proper cell types (Smolko et al., 2018; Ninova et al., 2020). A substantial fraction of the H3K9me-rich heterochromatin in which cryptic initiation concentrates (Fig. 3E) also carries H3K36me2 (Chaouch et al., 2021; Goto et al., 2024; Jayakrishnan et al., 2025), and on salivary gland polytenes, NSD protein is enriched in pericentric heterochromatin and across chromosome 4 (Lindehell et al., 2021). The regions at risk for cryptic initiation might therefore differ in how their H3K36me2 is maintained. Because the enzyme that supplies the mark at a given locus cannot be determined from steady-state profiling, we took advantage of S2-cell profiles of H3K36me2 and H3K36me3 following RNAi depletion of Set2, NSD, or both (Jayakrishnan et al., 2025), from which the compartments below are derived.

Viewed genome-wide, regions carrying both H3K9me2/3 and H3K36me2 occupy the pericentromeric portion of every major arm and the whole of chromosome 4 (Fig. 6A). This territory contains the bulk of the transposons and the major piRNA clusters, and the upregulated extragenic cryptic sites fall preferentially inside it. Across the major arms, 11.9 Mb carries both H3K9me and H3K36me2 (Fig. 6B). H3K36me3 is almost entirely nested within this compartment: of the 6.1 Mb of H3K9me chromatin that carries K36me3, 5.8 Mb also carries K36me2. The dual HC compartment is therefore defined by H3K9me together with H3K36me2, and H3K36me3 is a subset within it rather than an independent state.

**Figure 6.**
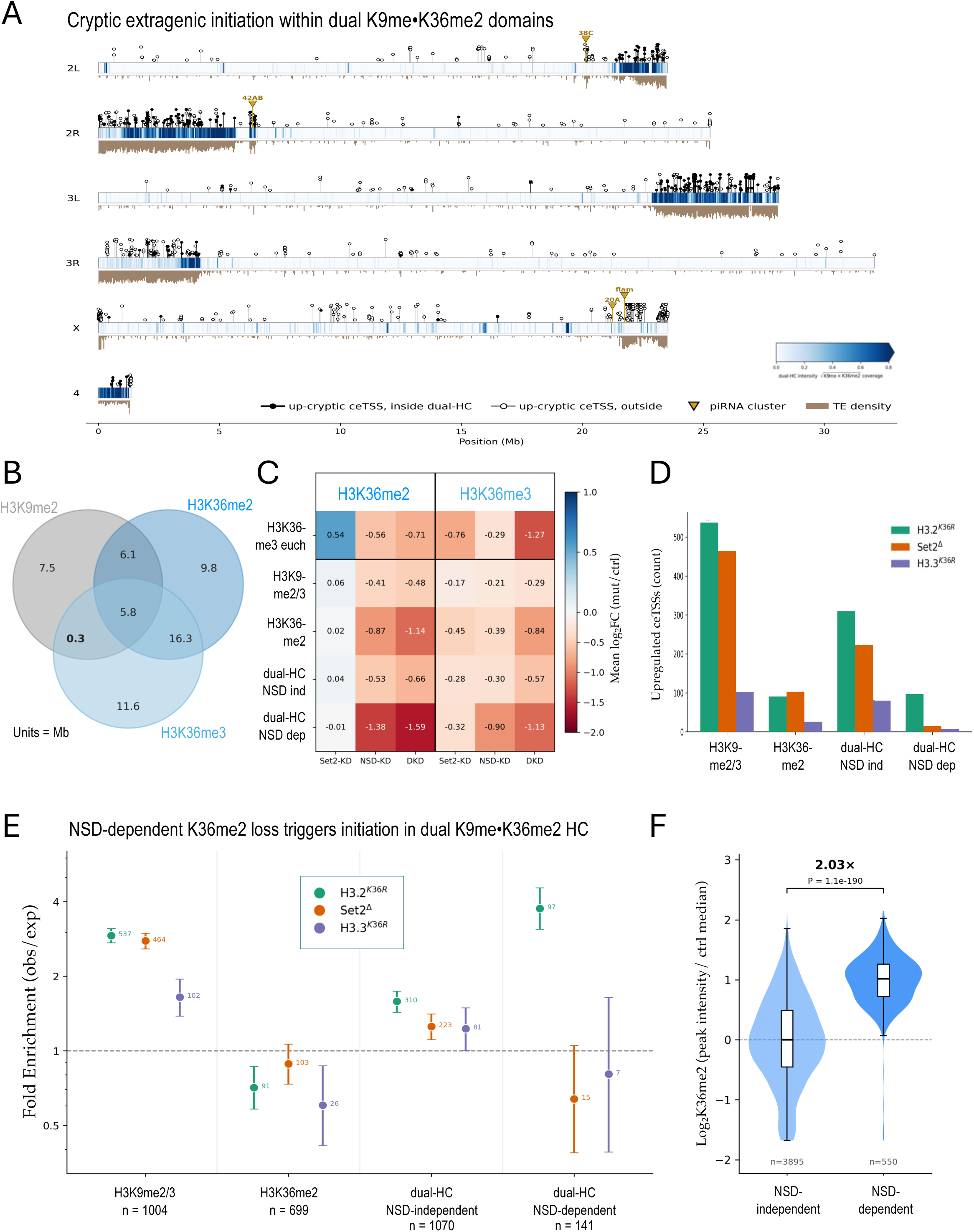
NSD-dependent H3K36me2 loss triggers initiation in dual K9me•K36me2 heterochromatin. **(A)** Genome-wide view of upregulated cryptic extragenic TSSs (lollipops) overlaid on dual HC (defined in panel B). Filled lollipops are inside dual HC; open ones are outside. Transposon density and annotated piRNA clusters are also shown. **(B)** Genomic overlap (in Mb) between H3K9me2/3, H3K36me2 and H3K36me3 across the major chromosome arms in S2 cells (Jayakrishnan et al., 2025). **(C)** Change in H3K36me2 and H3K36me3 levels (mean log₂ fold change) following RNAi of Set2 (Set2-KD), NSD (NSD-KD), or both (DKD) in S2 cells, across five compartments: a euchromatic H3K36me3 control, H3K9-only, H3K36me2-only, and NSD-independent and NSD-dependent dual HC. **(D, E)** Upregulated extragenic cryptic sites in four compartments, shown as raw counts (D) or fold-enrichment (E) over the tested background, for each genotype. **(F)** H3K36me2 peak intensity (log2, normalized to the NSD-independent median) at peaks in NSD-independent or NSD-dependent dual HC.

To determine which enzymes maintain H3K36me2 within this compartment, we scored H3K36me2 and H3K36me3 changes after knockdown of Set2 (Set2-KD), NSD (NSD-KD), or both (DKD) (Fig. 6C; Methods). In a euchromatic control compartment, Set2 knockdown raised H3K36me2 levels (+0.54) and lowered those of H3K36me3 (−0.76), the expected signature of a blocked conversion from di- to trimethylation. Within dual HC, NSD knockdown produced a loss of H3K36me2 over 2.3 Mb (NSD-dependent dual HC), whereas the remaining 9.9 Mb showed no significant loss (NSD-independent). Set2 knockdown had no effect on H3K36me2 in either subcompartment, whereas NSD knockdown reduced it strongly in the NSD-dependent one (−1.38; −1.59 on double knockdown) and more modestly in the NSD-independent one (−0.53). Set2 therefore makes no measurable contribution to H3K36me2 anywhere in dual HC; the two domains are distinguished by their degree of NSD dependence.

Mapping the upregulated extragenic cryptic sites onto these compartments separates the genotypes (Figs. 6D,E). All three genotypes were enriched in H3K9-only chromatin (*H3.2^K36R^*, 2.9-fold; *Set2Δ*, 2.8-fold; *H3.3^K36R^*, 1.6-fold), and none was enriched in K36me2-only chromatin that lacks H3K9me (0.60- to 0.89-fold). In NSD-independent dual HC all three mutants were modestly enriched (1.2- to 1.6-fold). Only in the NSD-dependent fraction of dual HC do the genotypes diverge: *H3.2^K36R^* rose to 3.8-fold (97 of 141 sites), whereas *Set2Δ* fell to 0.64-fold and *H3.3^K36R^* to 0.81-fold.

Within dual HC, therefore, the regions wherein NSD is the sole enzyme supplying H3K36me2 are the ones that distinguish the three mutant genotypes. Removing the methyl-accepting residue derepresses cryptic initiation at these loci; removing Set2 does not. The distinction is not a positional one: NSD-dependent and NSD-independent dual HC are interleaved along the chromosomes and both carry H3K9me and H3K36me2. Nor is it a matter of how much dimethyl mark is present. In control cells, H3K36me2 peak intensity in the NSD-dependent fraction is 2.0-fold higher than in the NSD-independent fraction (P = 1.1 × 10⁻¹⁹⁰; Fig. 6F). What separates these two fractions is the enzyme that deposits the dimethyl mark, a property revealed only by genetic perturbation. Thus, H3K36me2 itself, and not the trimethyl product of Set2, is the mark that protects these loci from cryptic initiation.

## Discussion

Large regions of metazoan genomes are transcribed at either low or undetectable levels in somatic cells. Transposons, satellites and other repetitive sequences, along with the entire germline program, are present in every nucleus and essentially stay silent during the life of the animal. The results presented here identify H3K36me2 as one of the marks that enforces this silence (Chaouch et al., 2021; Lindehell et al., 2023; Padilla et al., 2025), and place its requirement at the step of initiation rather than at elongation or RNA turnover. *Set2*Δ deletes a single writer and leaves the substrate intact, whereas the K36R mutants leave every writer intact and remove the substrate from one H3 variant. Comparing the two resolves which methylation state is required at a locus rather than which enzyme delivers it, a distinction that enzyme knockouts cannot make and that could not be addressed in the dual K9•K36 HC first described in mouse embryonic stem cells (Barral et al., 2022).

### Mechanistic Control of Cryptic Initiation

In budding yeast, Set2 and H3K36me suppress initiation from cryptic promoters within transcribed gene bodies, coupled to histone deacetylation in the wake of elongating polymerase. In the fly, intragenic cryptic initiation affects a small (∼1%) fraction of testable sites in any genotype, and the sites that do respond are uncoupled from host-gene transcription. Loss of K36 methylation perturbs acetylation over the 3′ ends of K36me3-rich genes (Figs. 2G; S2H,I), but it does not translate into widespread intragenic initiation. Although RPD3S deacetylase almost certainly still operates over H3K36me3-rich chromatin in flies, its role in suppressing cryptic initiation is not supported by our data. *H3.3^K36R^* makes the point most sharply, activating only 47 intragenic sites against 638 extragenic ones. Concurrent work in human cells is consistent with this view: loss of SETD2 catalytic activity produced cryptic initiation largely near the 3′ ends of a subset of genes, coupled to defective termination, and attributed to indirect consequences of SETD2 activity rather than to loss of H3K36me3 itself (Kopczyńska et al., 2026). We conclude that the yeast configuration is a derived condition rather than the ancestral one. *S. cerevisiae* lost H3K9 methylation, HP1 and the piRNA pathway from a more complex ancestor, and it carries a small, largely inactive transposon burden. Thus the yeast anti-cryptic function for K36me3 may have evolved after dual HC was lost.

Transposons and repeats are almost always scored by counting the RNA that accumulates over them, a measurement that sums synthesis, processing and decay and cannot locate the site at which transcription began. Initiation-level data are not simply a more sensitive version of the same measurement. A nascent TSS is a coordinate, which can be intersected with a chromatin compartment or placed inside or outside a gene body; a family-level fold change supports neither operation, and multi-mapping makes locus assignment unreliable at the RNA level in any case. The distinction matters most because the surveillance machinery is itself a target of the lesion. Loss of K36 methylation activates piRNP biogenesis factors in somatic tissue (Fig. 5), so RNA-level measurements of repeat expression in these animals reflect the balance between new synthesis and piRNA-directed decay. Indeed, reducing *krimper* in *H3.3^K36M^* eye discs unmasked element activation that surveillance had been concealing (Chaouch et al., 2021). Our own polyA RNA-seq data therefore understate the initiation defect, and the two readouts should not be treated as interchangeable.

Dual HC, the compartment defined here in Figures 3, 4 and 6, has been described independently in several other systems. These domains were first identified in mouse embryonic stem cells as thousands of loci dependent on SETDB1 and on NSD enzymes rather than on SETD2 (Barral et al., 2022). Removing H3K36 methylation in mouse mesenchymal stem cells redistributes H3K9me3 out of constitutive heterochromatin and collapses genome compartment structure (Padilla et al., 2025). In *Drosophila*, H3K36me2 marks pericentric transposons in the eye disc, and its redistribution accompanies transposon derepression (Chaouch et al., 2021); the mark is delivered to these loci via ISWI-dependent recruitment of NSD (Goto et al., 2024). In fly brain, loss of K36 methylation derepresses transposons and activates piRNA-pathway genes (Lindehell et al., 2023), and in the hindgut, linker histone H1 recruits NSD to establish H3K36me2 upstream of H3K9me3, restraining transposon mobilization (Luo et al., 2026b). None of this prior work mapped where transcription initiates. Our data supply that missing information layer, and it is precisely this layer that reveals a *transcriptional consequence*: the extragenic TSSs that fire aberrantly in *H3.2^K36R^* are enriched at loci that depend on NSD for H3K36me2. Note, we have not measured the NSD dependence of these sites in the wing disc. However, the two datasets were generated entirely independently, so their correspondence should be viewed as a cross-system result rather than a within-tissue one.

Within dual HC domains, the two K36 methylation states are not equivalent. *H3.2^K36R^* activates cryptic initiation in the NSD-dependent fraction 3.8-fold, whereas *Set2*Δ falls below expectation at the same loci. *Set2*Δ animals retain H3K36me2 in this compartment and lose only the trimethyl product, so the protective activity resides in the dimethyl state. The residue mutants are what make the conclusion available: an effect seen only on deleting *Set2* could reflect non-catalytic functions of the enzyme, whereas removing the methyl-accepting lysine leaves the enzymes intact and reports on the mark. The K36me2-to-me3 conversion and its regulation are under separate investigation. An independent line of work in fly somatic tissue has reached a parallel conclusion for the other residue: H3K9 dimethylation on H3.2 is required for transposon silencing in the hindgut, and H3K9me3 is not sufficient, since silencing fails even when trimethylation rises (Luo et al., 2026a). For both residues, the lower methylation state carries the requirement.

The two K36R genotypes are not equally penetrant, and the difference tracks the compartments the variants occupy rather than the severity of the mutations. In somatic tissue, H3.2 concentrates in DAPI-dense heterochromatin and colocalizes with H3K9 methylation, whereas H3.3 prefers euchromatin and telomeres (Mito et al., 2005; Schwartz and Ahmad, 2005; Lewis et al., 2010). Replacing K36 in the replication-dependent histones therefore removes the mark from the compartment where it is required, whereas replacing it in H3.3 largely does not. This interpretation accounts for the pattern across our data: *H3.2^K36R^* separates from the other genotypes specifically in NSD-dependent dual HC, and *H3.3^K36R^* shows no enrichment there.

### Implications for Genome Stability and Disease

Germline derepression is a separate consequence of the same lesion. Somatic wing cells lacking H3K36 methylation express germline-restricted genes and piRNP biogenesis factors, and 5′ PRO-seq shows that the promoters of those genes fire at the annotated start site, specifically in *H3.2^K36R^*. The derepression is therefore initiation-driven and local, rather than a consequence of transcripts escaping turnover. The affected genes are also not switched on as a unit, and the H3K9 methyltransferases are unchanged in every genotype. We therefore report two separable consequences of removing H3K36me, and leave the relationship between them open.

Whether cryptic transcription of transposons actually leads to their mobilization is a question our data cannot address, but it has been answered directly elsewhere. Long-read sequencing of fly somatic tissue in which heterochromatin assembly is compromised recovers new LTR retrotransposon insertions together with DNA damage, establishing that transcriptional derepression in this compartment carries a genomic cost (Luo et al., 2026b).

Three questions follow from this work. First, what factors read the dual HC mark? In transcribed genes, H3K36me3 recruits a deacetylase complex in the wake of elongating polymerase, a mechanism that cannot operate in a compartment that is not transcribed. Identifying the effector that converts H3K36me2 into silent heterochromatin would explain how one residue serves opposing functions on either side of the euchromatic boundary. Second, what is the order of events? Loss of SETDB1 removes both marks from dual domains in mouse ES cells (Barral et al., 2022), whereas transgenic expression of mutant histones in the fly hindgut places H3K36me2 upstream of H3K9me3 (Luo et al., 2026b), and silencing there depends on H3K9me2 rather than H3K9me3 (Luo et al., 2026a). Because steady-state profiles cannot establish order, resolving this question will require following H3K9 methylation states and HP1 occupancy through development after K36 methylation is lost. Third, does the same vulnerability extend to human disease? H3K36M and loss of NSD1 both deplete H3K36me2 in human tumors, so our results predict that these tumors should fail at cell identity as well as at suppressing transposition. Expression of germline and testis-restricted programs in existing tumor datasets offers a direct readout. If H3K36 dimethylation is the mark that restricts the germline program from somatic cells, then the oncohistone that removes this mark is likely doing more than simply disabling a growth-suppressive pathway.

## Methods

### Drosophila strains and husbandry

Set2Δ animals carry the Set2^1^ null allele together with its rescue transgene (Larschan et al., 2007; Bloomington Stock Center no. 77,917), and are compared to genotype-matched yw controls (Salzler et al., 2024).

*H3.2^K36R^* and *H3.2^HWT^* animals carry a 12x array of the 5 kb replication-dependent histone repeat unit, mutant (H3.2 lysine-36 to arginine) or wild type, integrated at the VK33 attP site and hemizygous over a null deletion of the endogenous histone locus, Df(2L)HisC^ED1429^ (*ΔHisC*) (McKay et al., 2015; Meers et al., 2017; Salzler et al., 2023).

*H3.3^K36R^* animals carry a CRISPR/Cas9-generated *H3.3B^K36R^* allele as the sole source of H3.3, on an *H3.3A* null background (*H3.3A^2x1^*; Sakai et al., 2009). *H3.3^Ctrl^* animals carry wild-type *H3.3B* on the same background (Salzler et al., 2023).

Crossing schemes and complete genotypes are given in McKay et al. 2015, Salzler et al. 2023, and Salzler et al. 2024. Larvae were picked at the L2 stage (50 per vial) from egg-laying cages and raised on standard cornmeal-molasses food at 25°C until the wandering third-instar (WL3) stage. Three biological replicates (each derived from an independent collection of larvae/vials) were processed per genotype for PRO-seq, RNA-seq, and CUT&RUN.

### Isolation of larval nuclei for PRO-seq

Wing imaginal discs (typically 100 per biological replicate) were dissected from wandering third-instar larvae into Grace’s insect medium, pelleted (800×g, 4°C, 5 min) and washed in nuclei extraction buffer (NEB; 10 mM HEPES-KOH, pH 7.5, 2.5 mM MgCl2, 10 mM KCl). Discs were homogenized in 1 mL NEB in a 2 mL Dounce homogenizer (20 strokes of the loose pestle, 10 min on ice, 15 strokes of the tight pestle), filtered through a 30 µm CellTrics filter, and nuclei were pelleted (1000×g, 4°C); all tips, glassware, and homogenizer components were pretreated with NEB containing 5% BSA. Nuclei were checked by DAPI staining, then permeabilized and cryopreserved at –80°C in Buffer F (50 mM Tris-Cl, pH 8.0, 40% [v/v] glycerol, 5 mM MgCl2, 1.1 mM EDTA, 0.5 mM DTT) with SUPERase-In RNase inhibitor.

### PRO-seq library construction and data analysis

For each sample, 1 million permeabilized nuclei were used for nuclear run-on, with human cells added for spike-in normalization. Nuclear run-on assays and library preparation were performed as described (Mahat et al., 2016; Mimoso and Goldman, 2023), and libraries were sequenced on the Illumina NovaSeq platform. Reads were processed with custom scripts available on the AdelmanLab GitHub (https://github.com/AdelmanLab/NIH_scripts). Unique Molecular Identifiers (UMIs) were extracted using UMI-tools (Smith et al., 2017), adapters were trimmed using cutadapt 1.14 (Martin, 2011), and paired-end reads were mapped to a combined hg38 (spike-in) and dm6 genome index using bowtie2 (Langmead and Salzberg, 2012). Properly paired reads were separated by genome, deduplicated using UMI-tools, sorted with samtools (Li et al., 2009) and converted to bedGraphs at single-nucleotide resolution. Samples displayed comparable recovery of spike-in reads; PRO-seq samples were therefore normalized using size factors from DESeq2 (Love et al., 2014). Replicate bedGraphs were merged within conditions and depth-normalized to generate bigWig files binned at 1 bp, 10 bp or 25 bp intervals depending on the downstream analysis.

### RNA-seq library preparation and analysis

RNA from 25–30 wing imaginal discs per biological replicate was extracted with TRIzol and purified with the Zymo RNA Clean & Concentrator-5 kit, including DNase I treatment. Poly(A)-enriched, strand-specific libraries (KAPA Stranded mRNA-Seq Kit, Roche) were sequenced on an Illumina NovaSeq S4 platform using 50-bp paired-ends.

Reads were processed using the nf-core/rnaseq pipeline (v3.17.0; Ewels et al., 2020), implemented in Nextflow (Di Tommaso et al., 2017): reads were trimmed using Trim Galore, aligned to dm6 using STAR 2.7.11b (Dobin et al., 2013) in two-pass mode with -- outFilterMultimapNmax 20, and quantified using Salmon 1.10.3 (Patro et al., 2017), with quality control summarized by MultiQC (Ewels et al., 2016). Transcript-level estimates were summarized to genes using tximport (Soneson et al., 2015).

### CUT&RUN library construction and data analysis

CUT&RUN against pan-acetylated histone H4 (H4ac) was performed on dissociated wing imaginal discs from wandering third-instar larvae (20 discs per replicate, biological triplicate), as described (Skene and Henikoff, 2017; Ahmad and Spens, 2019). Paired-end reads were aligned to dm6 using bowtie2 in local mode, filtered for MAPQ ≥ 5, and deduplicated using Picard MarkDuplicates. After quality control, three replicates were retained for *Set2*Δ, *H3.2^K36R^* and H3.2HWT, and two for *yw*, *H3.3^K36R^* and *H3.3^Ctrl^* (Fig. S2A–F). A *Drosophila yakuba* spike-in was recovered inconsistently and was not used. Alignments were instead converted to RPGC-normalized bigWig files, and TMM normalization factors computed from the CUT&RUN counts were close to 1.0 for all libraries, consistent with the absence of a global change in H4ac by western blotting.

### Refinement of gene annotation (GGA) using PRO-seq and RNA-seq

To pair gene-level features with PRO-seq data, a dominant TSS (domTSS) and transcription end site (TES) were assigned to each active gene using GGA (https://github.com/AdelmanLab/GetGeneAnnotation_GGA), which combines RNA-seq read abundance (HISAT2 v2.2.1 alignments; Kim et al., 2019) with PRO-seq R2 reads (RNA 5’ ends) from all conditions. Fly-specific parameters, established in consultation with the method’s developers, were a promoter-distal cutoff of 200 bp and an annotation join distance of ∼150–200 bp (500 bp default).

TSScall (Scruggs et al., 2015) was applied to the combined PRO-seq dataset across all eighteen libraries. Called events falling within ±100 bp of an annotated FlyBase (dm6 r6.32) gene start were classified as observed TSSs (obsTSS); the single most highly used TSS per gene was designated as the domTSS, and additional called sites at the same gene as annotated alternative TSSs (aaTSS). All remaining events were classified as novel unannotated TSSs (nuTSS) and subdivided by position into potential intragenic (piTSS) and potential extragenic (peTSS) TSSs. For Figure 1D, 5’ PRO-seq reads within ±75 nt of each called position were summed across all eighteen libraries and binned into 80 equal-width log10 intervals.

### H4ac meta-plots

Genes were ranked by H3K36me3 over the gene body in control S2 cells (Jayakrishnan et al., 2025), and high- and low-K36me3 sets were taken as the top and bottom quartiles, with the low set restricted to genes expressed in S2 cells. All gene sets were filtered for expression in wing discs (mean TPM ≥ 1 in yw and H3.2^HWT^), location on the major chromosome arms and length ≥ 1.5 kb, excluding *mod(mdg4)*. RPGC-normalized H4ac signal (replicates pooled) was profiled with deepTools computeMatrix (Ramírez et al., 2016) over gene bodies scaled to 5 kb with 2 kb flanks, in 25-bp bins; RPGC normalization equalizes genome-wide coverage, so differences reflect redistribution of H4ac rather than changes in its total amount. The per-gene log2 ratio (mutant/control) over the 3′ half of the gene body was regressed on K36me3, with control H4ac level and S2 expression as covariates (n = 5,346 genes).

### Differential expression analysis

For gene-level PRO-seq analysis, reads were summed within the TSS-to-TES window for each active gene using make_heatmap (https://github.com/AdelmanLab/NIH_scripts) and tested with DESeq2 (Wald test; Love et al., 2014).

Novel TSSs were filtered for expression with edgeR filterByExpr, applied separately to piTSS and peTSS, yielding 22,156 candidate cryptic intragenic TSSs (ciTSS) and 8,632 candidate cryptic extragenic TSSs (ceTSS). Because Drosophila TSSs are predominantly narrow (Meers et al., 2018b), reads were summed in a 21 bp window (±10 bp) centered on each called TSS. Each mutant was compared to its matched control (*Set2Δ vs. yw; H3.2^K36R^ vs. H3.2^HWT^; H3.3^K36R^ vs. H3.3^Ctrl^*) using the quasi-likelihood framework in edgeR (Robinson et al., 2010).

### Chromatin state, repeat element, and piRNA cluster enrichment analyses

Upregulated ciTSS and ceTSS (foreground), and all other tested sites of the same class in that comparison (background), were assigned to the nine chromatin states defined in BG3 cells (Kharchenko et al., 2011), using the ±10 bp window and one state per site, with unmapped regions designated state 0 (Fig. 3D). For Fig. 3E, unmapped sites were removed and enrichment was calculated as log2[(observed + 0.5)/(expected + 0.5)], where the expected value is the foreground total multiplied by the fraction of background sites in that state. Significance was assessed by two-sided Fisher’s exact test without correction; Benjamini-Hochberg-adjusted values were used, and no state changed significance after adjustment.

Repeat element overlap was assessed by intersecting the same ±10 bp windows, irrespective of strand, with dm6 RepeatMasker annotations from the UCSC Genome Browser; a site could be scored in more than one class, and satellite subfamilies were pooled. For each genotype, upregulated ciTSSs and ceTSSs were pooled and compared with all 30,788 tested sites; the expected count for a class is the number of upregulated sites multiplied by the fraction of tested sites in that class. Significance was assessed by two-sided Fisher’s exact test with Benjamini-Hochberg correction across the twelve genotype-by-class comparisons (Fig. 4D). PRO-seq alignments retain multi-mapping reads (bowtie2 defaults, MAPQ 0–1), and 54.4% of all ceTSS fall within repetitive or transposon-derived sequence, a fraction corroborated by an empirical mappability filter; consequences for locus-level assignment are considered in the Results.

piRNA cluster coordinates (Chaouch et al., 2021) were intersected with peTSSs. Fold-enrichment at each cluster was calculated as observed over expected peTSS overlap given cluster size, and significance was assessed by Fisher’s exact test.

### Dual heterochromatin (dual HC) compartment definition and NSD dependence

Compartments were defined from spike-in-normalized ChIP-seq of H3K36me2, H3K36me3, and H3K9me2 in S2 cells after RNAi of Set2, NSD, or both (DKD), with GFP-RNAi controls (Jayakrishnan et al., 2025; GSE253391). Control-cell peak calls defined four mutually exclusive compartments: dual heterochromatin (dual HC; H3K9me2 together with H3K36me2 and/or H3K36me3, 12.2 Mb, of which 11.9 Mb carries H3K36me2), H3K9me2-only (7.55 Mb), H3K36me2-only (9.78 Mb), and a euchromatic H3K36me3 reference territory (H3K36me3 peaks outside H3K9me2 and dual HC, 27.96 Mb). The portions of dual HC overlapping a set of intervals with ≥4-fold loss of H3K36me2 upon NSD depletion (Jayakrishnan et al., 2025; overlapping regions merged) were designated NSD-dependent (2.31 Mb, 746 intervals) and the remainder NSD-independent (9.87 Mb, 4,473 intervals).

For Fig. 6A, dual HC intensity was scored in 150-bp bins as the geometric mean of S2 H3K9me and H3K36me2 coverage, smoothed and averaged into 40-kb windows. For Fig. 6C, H3K36me2 and H3K36me3 signal (three replicates per condition) was extracted for each compartment using deepTools multiBigwigSummary (Ramírez et al., 2016), averaged across bins within each interval and then across intervals, and expressed as log2[(condition signal + 0.05)/(control signal + 0.05)].

For Figs. 6D and E, analysis was restricted to the 7,768 ceTSSs testable in all three comparisons, excluding 864 chrY sites because the S2 compartment reference is undefined on chrY. Each ceTSS was assigned to a compartment by the position of its TSS: H3K9me2-only (n = 1,004), H3K36me2-only (n = 699), NSD-independent dual HC (n = 1,070) and NSD-dependent dual HC (n = 141). Fold-enrichment was calculated as the fraction of a genotype’s upregulated ceTSSs falling in a compartment divided by the fraction of all tested ceTSSs in that compartment, with Wilson 95% confidence intervals and a one-tailed hypergeometric test. The four compartments are a selection rather than a partition of the genome.

For Fig. 6F, MACS2 bdgpeakcall scores (spike-in-scaled, input-normalized ChIP signal; Jayakrishnan et al., 2025) of control H3K36me2 peaks overlapping NSD-dependent (n = 550) and NSD-independent (n = 3,895) dual HC were normalized to the NSD-independent median, log2-transformed, and compared by two-sided Mann-Whitney U test; the fold difference is the ratio of medians.

### Cross-study and first-instar gene expression comparisons

Differentially expressed genes were defined as padj < 0.05 and |log2 fold change| > 1 (DESeq2). For Fig. 5B, upregulated genes from the three wing-disc comparisons were intersected with DESeq2 results for H3.3^K36M^ expressed under ey-GAL4 control in the eye-antennal disc, compared with *yw* (Chaouch et al., 2021), among the 11,080 genes tested in every dataset. First-instar (L1) whole-larval poly(A)+ RNA-seq data for H3.2^K36R^ and the combined *H3.3^K36R^;H3.2^K36R^* mutant were generated here (GEO accession GSE196114) and analyzed in the same way (Fig. 5D). Enrichment of germline-pathway genes among L1 upregulated genes was assessed by one-tailed hypergeometric test against the 25 Fig. 5C genes tested in the L1 data, including the H3K9 methyltransferase controls.

### Nascent transcription at germline gene dominant TSSs

For each mutant/control pair, domTSSs with ≤10 pooled 5′ PRO-seq reads in the matched control were classed as silent (n = 933 for H3.2^HWT^, 1,034 for yw and 944 for the H3.3 control). Change at each domTSS was expressed as log2[(mutant + 4)/(control + 4)]; a gain from 0 to 12 reads scores exactly 2.0, the fourfold threshold used in the text. Within each silent class, the germline and piRNA-pathway genes shown in Fig. 5C were compared with all other class members by two-sided Fisher’s exact test (proportion gaining ≥4-fold) and two-sided Mann-Whitney U test (distributions of log2 ratios). Because 21 of the 29 genes in Fig. 5C fall below the edgeR expression filter, per-gene significance is reported only for the eight that were tested.

### Statistical tests

Tests are described in the sections above; Benjamini-Hochberg correction was applied where indicated. Where a single correlation is reported across the three genotype comparisons (Fig. 2C), per-comparison coefficients were combined using Fisher’s z-transformation.

### Western blotting

Lysates of two wandering third-instar larvae per sample were prepared in SUTEB buffer (1% SDS, 8 M urea, 10 mM Tris-Cl pH 8.0, 10 mM EDTA pH 8.0) with β-mercaptoethanol, sodium butyrate and protease inhibitors, sonicated and clarified by centrifugation. Equal masses of protein (20 or 40 µg; Qubit Protein Assay) were resolved on 4–15% Mini-PROTEAN TGX Stain-Free gels (Bio-Rad), transferred to nitrocellulose and probed with rabbit anti-H4ac (Active Motif, #39925, 1:1000) and rabbit anti-H3K36me3 (Abcam, ab9050, 1:2000), with rabbit anti-H3 (Abcam, ab1791, 1:60000) and anti-H4 (GenuIN Biotech, #2134, 1:5000) as loading controls and goat anti-rabbit IgG-HRP (Sigma-Aldrich, #12-348, 1:5000) as secondary antibody; blots were stripped (Restore, Thermo Fisher Scientific) between probes. Band intensities were quantified by densitometry, and the H4ac/H3pan and H3K36me3/H4pan ratios for each mutant were normalized to its matched control and compared by two-tailed paired t-test on log2 fold change (Fig. S2G).

## Acknowledgements

We thank A. Goli for technical assistance. This work was supported by the National Institutes of Health (NIGMS) grant R35-GM136435 (to AGM).

---

**Figure S1.**
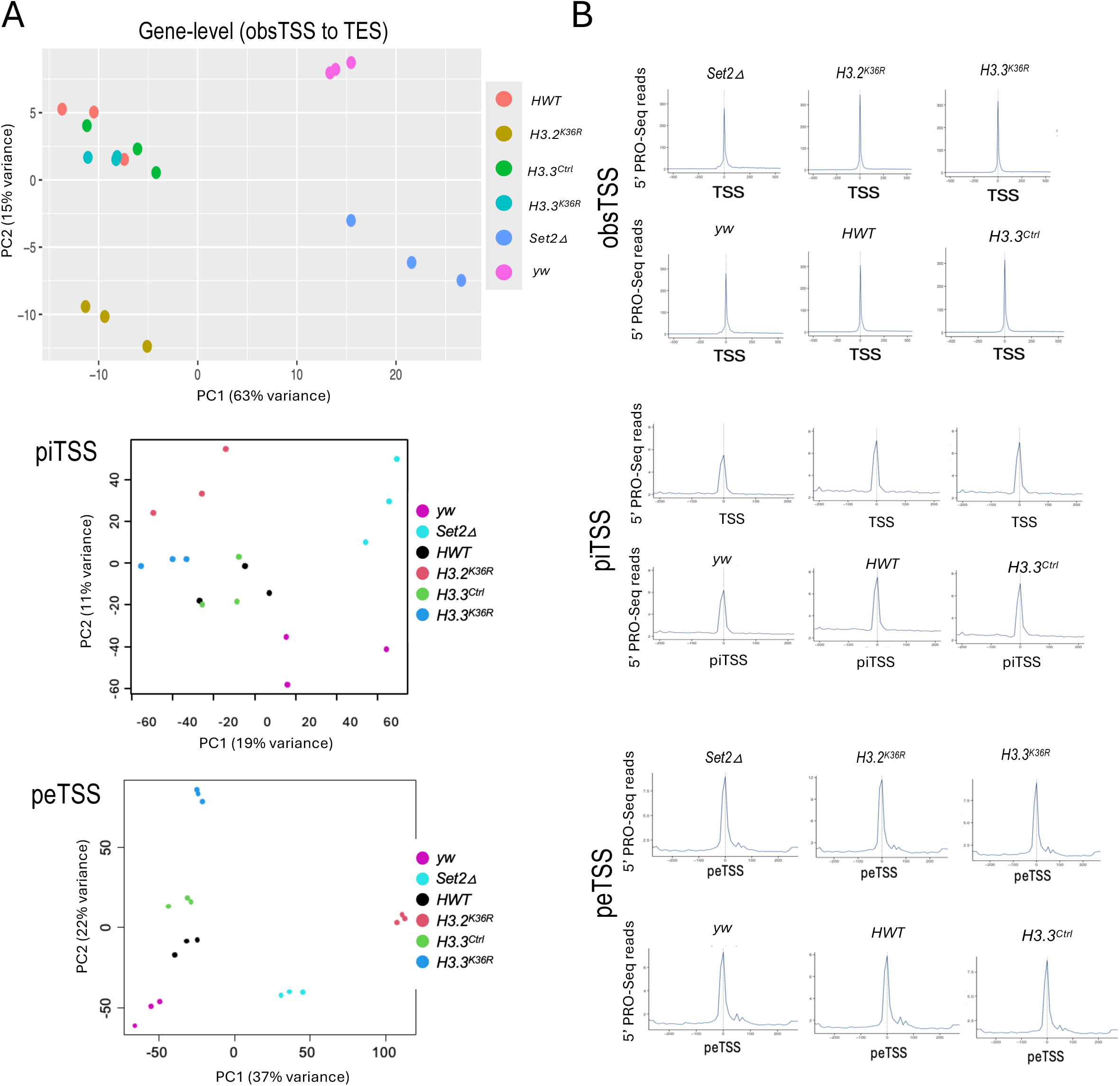
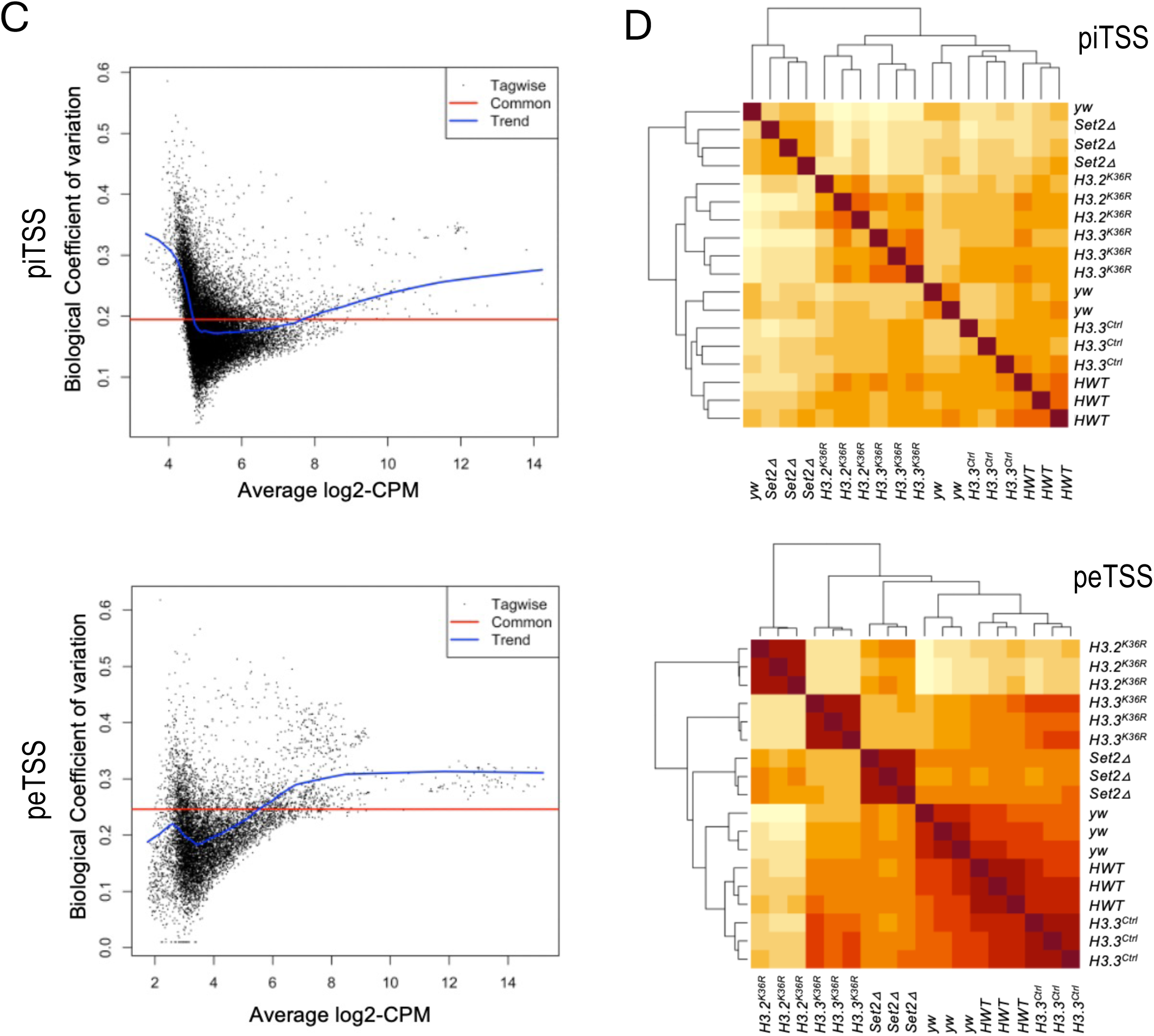
Quality control of PRO-seq libraries across genotypes. (A) Principal component analysis (PCA) of PRO-seq counts at three levels: gene level (reads summed from the observed TSS to the TES of each gene; top), potential intragenic TSSs (piTSSs; middle) and potential extragenic TSSs (peTSSs; bottom). Gene-level counts were DESeq2-normalized and rlog-transformed; piTSS and peTSS counts were taken in 21-bp windows centered on each called TSS (TSS ± 10 bp; Methods). Each point is one biological replicate. Replicates cluster by genotype at the gene and peTSS levels; piTSS counts, which are lower, separate genotypes less cleanly. (B) Meta-plots of 5′ PRO-seq signal centered on obsTSSs (top; ±500 nt), piTSSs (middle; ±200 nt) and peTSSs (bottom; ±200 nt). For each TSS class, mutants are shown above their matched controls. (C) Biological coefficient of variation (BCV) plotted against average abundance (log2 CPM) for piTSSs (top) and peTSSs (bottom). The red line marks the common dispersion across all sites and the blue curve the abundance-dependent trend. (D) Hierarchical clustering of Pearson correlation coefficients between replicates for piTSS (top) and peTSS (bottom) counts.

**Figure S2.**
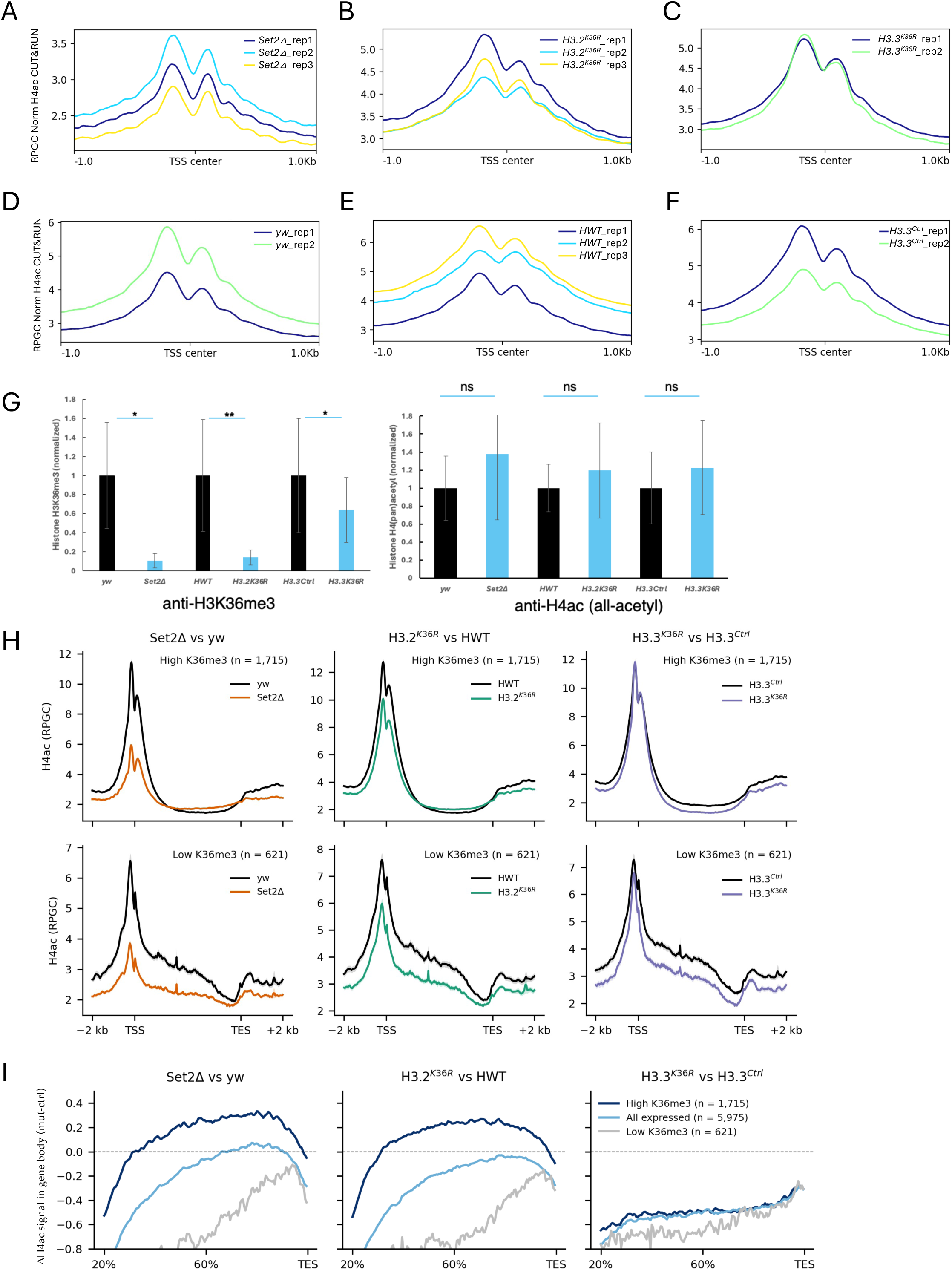
Histone H4 acetylation and H3K36me3 across genotypes. (A–F) RPGC-normalized H4ac CUT&RUN signal within ±1 kb of each gene’s annotated TSS, plotted separately for each biological replicate. Mutants are on the top row: Set2Δ (A), H3.2^K36R^ (B) and H3.3^K36R^ (C). Their matched controls are directly below: yw (D), HWT (E) and H3.3^Ctrl^(F). Replicates differ in overall signal level but share the same profile shape. (G) Quantification of western blots of whole-larval lysates for H3K36me3 (left) and H4ac (right). Band intensities were normalized to the loading control and expressed relative to the mean of the matched control; bars show mean ± SEM of three biological replicates. Significance by two-tailed paired t-test on log2 fold change (*, p < 0.05; **, p < 0.01; ns, not significant). (H) Meta-plots of H4ac CUT&RUN signal (RPGC, pooled replicates) across gene bodies scaled to 5 kb with 2 kb flanks, for genes with high (top; n = 1,715) or low (bottom; n = 621) K36me3. Controls are shown in black and mutants in color; shading, SEM across genes. Note that the reduced promoter signal in the mutants is common to all gene sets. (I) Difference in H4ac (mutant − control) from 20% of gene length to the TES, for genes with high K36me3, all expressed genes and genes with low K36me3. Gene sets are defined in Methods.

**Figure S3.**
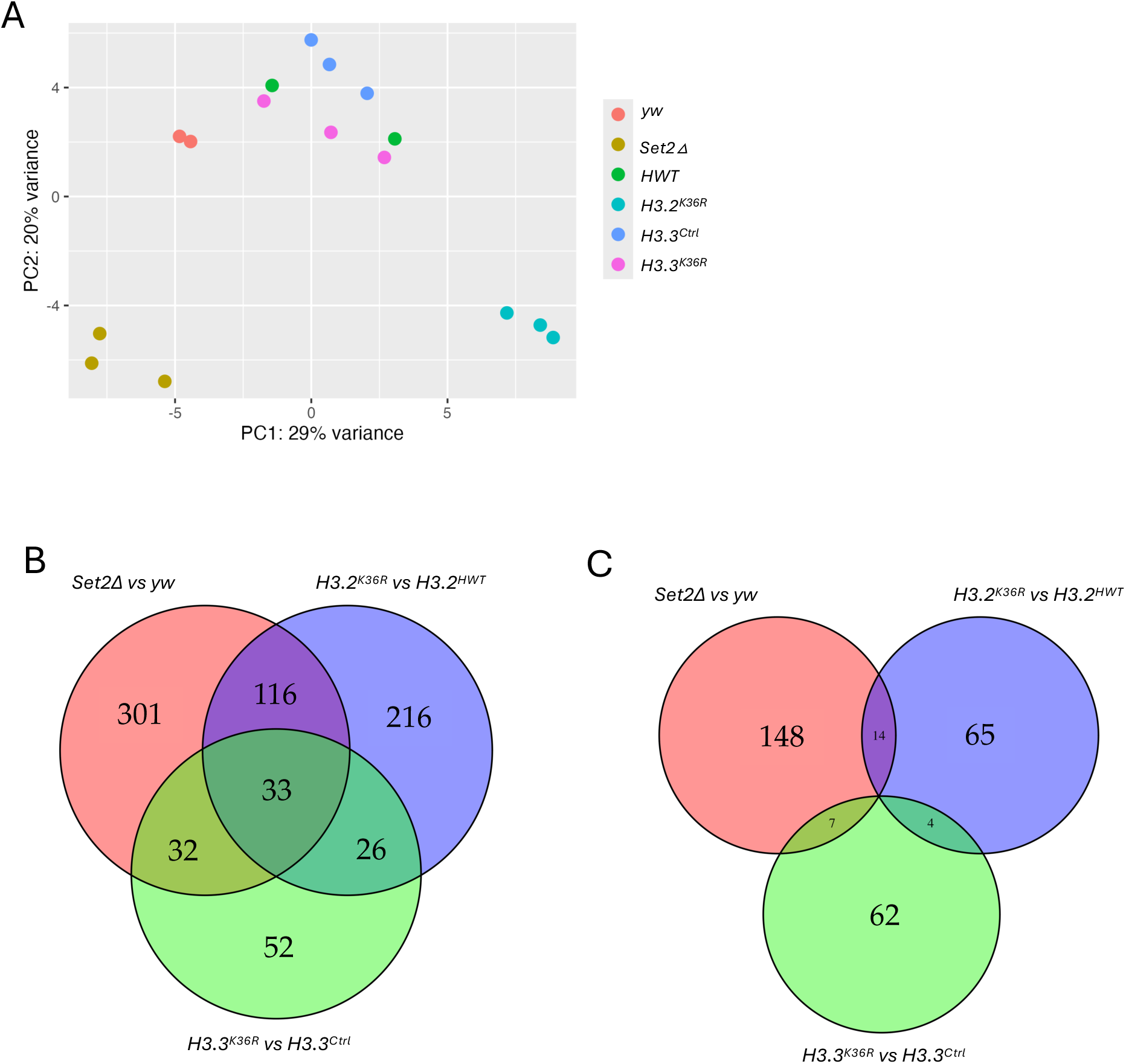
Polyadenylated RNA-seq: sample clustering and overlap of differentially expressed genes. (A) PCA of gene-level RNA-seq counts for the six genotypes; each point is one biological replicate. Replicates cluster by genotype, and Set2Δ and H3.2^K36R^ separate most clearly from the other samples. (B, C) Overlap among genes significantly upregulated (B) or downregulated (C) in Set2Δ vs. yw, H3.2^K36R^ vs. HWT and H3.3^K36R^ vs. H3.3^Ctrl^ (padj < 0.05, |log2FC| > 1).

